# Soluble TREM2 is a biomarker but not a mediator of fibrosing steatohepatitis

**DOI:** 10.1101/2025.03.11.642665

**Authors:** Joseph L. Dempsey, Christopher Savard, Vishal Kothari, Jingjing Tang, Sum P. Lee, Karin E. Bornfeldt, Rotonya M. Carr, George N. Ioannou

## Abstract

**Background:** Triggering receptor expressed on myeloid cells 2 (TREM2), a transmembrane, lipid-sensing protein expressed by Kupffer cells, is thought to play a role in metabolic dysfunction-associated steatotic liver disease (MASLD) and metabolic dysfunction-associated steatohepatitis (MASH). Plasma levels of the TREM2 cleavage product, soluble TREM2 (sTREM2), are strongly associated with MASLD severity. We investigated the role of TREM2 in MASH pathogenesis and whether sTREM2 acts both as biomarker and mediator of MASH.

**Methods:** Adult C57BL/6J mice were assigned to normal, high-fat (HF), or high-fat and high-cholesterol (HFHC) diets for 15, 30, 90, and 180 days. Plasma sTrem2 levels, liver pathology, and hepatic RNA expression were assessed. To test whether sTrem2 is a mediator of MASH, C57BL/6J mice were injected retro-orbitally with a liver-targeted adeno-associated virus, TBG-AAV8-sTrem2, resulting in secretion of sTrem2 by hepatocytes, or empty TBG-AAV8-control, and subsequently fed HFHC diet for 15, 90, or 180 days.

**Results:** HFHC-fed mice developed fibrosing steatohepatitis at 180 days together with a 15-fold increase in plasma sTrem2 levels. In the livers of HFHC-fed mice, crown-like structures (CLSs) consisting of TREM2^+^ macrophages surrounded necrotic, steatotic hepatocytes and their remnant lipid droplets, which contained prominent crystals containing cholesterol. TBG-AAV8-sTrem2-injected mice had higher levels of plasma sTrem2 than TBG-AAV8-control-injected mice, but there were no differences in liver weight, body weight, hepatic fibrosis, hepatic inflammation, hepatic cholesterol crystals, or plasma cholesterol levels.

**Conclusion:** TREM2^+^ macrophages characterize the CLSs that surround necrotic hepatocytes and their remnant lipid droplets and cholesterol crystals in fibrosing steatohepatitis-MASH. Plasma sTrem2 is a biomarker, but not a causal mediator, of MASH.

## INTRODUCTION

Triggering receptor expressed on myeloid cells 2 (TREM2) is a transmembrane, cell-surface receptor protein belonging to the superimmunoglobulin receptor family, which is expressed hepatically on Kupffer cells [1]. TREM2 can be divided into three major domains: 1) the N-terminus immunoglobulin domain, 2) the transmembrane domain, and 3) the short, C-terminus cytoplasmic domain. The TREM2 immunoglobulin domain can bind to phospholipids, lipoproteins, sulfatides, and lipopolysaccharides [2-8]. The TREM2 transmembrane domain interacts with the transmembrane adaptor proteins DNAX-activating protein 12 (DAP12) and DAP10 for intracellular signal transduction [9]. TREM2 ligand binding is associated with activation of several pathways including mitogen-activated protein kinase (MAPK), nuclear factor-kappa B (NF-κB), mechanistic target of rapamycin (mTOR), and actin cytoskeleton rearrangement [8, 10]. TREM2 activation may also competitively inhibit Toll-like receptor (TLR) signaling of extracellular signal-regulated kinase (ERK) by DAP12 recruitment of adapter proteins GRB2 and SOS [3, 8].

In the liver, TREM2^+^ macrophages recognize exposed phospholipids and cholesterol from apoptotic, steatotic hepatocytes, leading to their phagocytosis [11]. Progression of metabolic dysfunction-associated steatotic liver disease (MASLD) to fibrosing steatohepatitis (metabolic dysfunction-associated steatohepatitis; MASH) is characterized by the appearance of a subset of TREM2-expressing macrophages in the liver that are thought to be monocyte-derived Kupffer cells [11]. Initially identified in 2019, TREM2^+^ macrophages were found in proximity to fibrosis, and they were labeled as scar-associated macrophages (SAMac) [12]. Subsequently, they were also noted to express Cd63, Cd9, and Gpmnb in addition to Trem2, which are markers ascribed to “lipid-associated macrophages (LAM)” in other tissues [13]. Most research since has suggested that LAMs act to regulate and limit inflammation and fibrosis, remove necrotic cells, and have a protective effect overall [14, 15]. We have previously shown that macrophages aggregate around and process steatotic, necrotic hepatocytes that contain cholesterol crystals in their remnant lipid droplets, forming crown-like structures (CLSs) [16-20]. It was unknown if the macrophages that form CLSs are Trem2^+^.

In steatotic liver disease, the membrane-bound TREM2 (mTREM2) is cleaved by ADAM17 to shed the extracellular immunoglobulin domain as the soluble form of TREM2 (sTrem2; **Figure 1A**) [15, 21, 22], which can be measured in plasma. Cleavage of TREM2 (“ectodomain shedding”) can inactivate its downstream signaling function and disrupt its ability to interact with its ligands, which are critical for Kupffer cell activation. Plasma levels of sTREM2 are elevated in humans and mice with MASH and cirrhosis [14, 23-25], and sTREM2 is strongly associated with histological features of MASH with liver stiffness, suggesting it may be a non-invasive diagnostic biomarker [23]. Plasma sTREM2 has been associated with the recruitment and expansion of TREM2^+^ macrophages to fibrotic regions to limit MASH [14]. However, weekly injections of recombinant sTrem2 failed to protect Trem2-deficient mice from fibrosis after 8 weeks of diet-induced MASH [15]. Furthermore, bone marrow derived macrophages from wild-type mice co-cultured with AML12 cells (a mouse hepatocyte cell line) and treated with palmitic acid and sTrem2 had significantly decreased phagocytosis, suggesting a relationship between Trem2, sTrem2, and resolution of fibrotic regions in MASH [15]. In mouse primary hepatocytes, we demonstrated that sTrem2 chimeric protein and palmitate increased lipid droplet formation and triglyceride content [25]. While plasma sTREM2 levels appear to be a novel and strong biomarker of MASH severity in humans, it is unclear whether sTREM2 directly mediates any pathophysiological effects or merely reflects the aggregation of TREM2^+^ Kupffer cells that characterize MASH.

**Figure 1.**
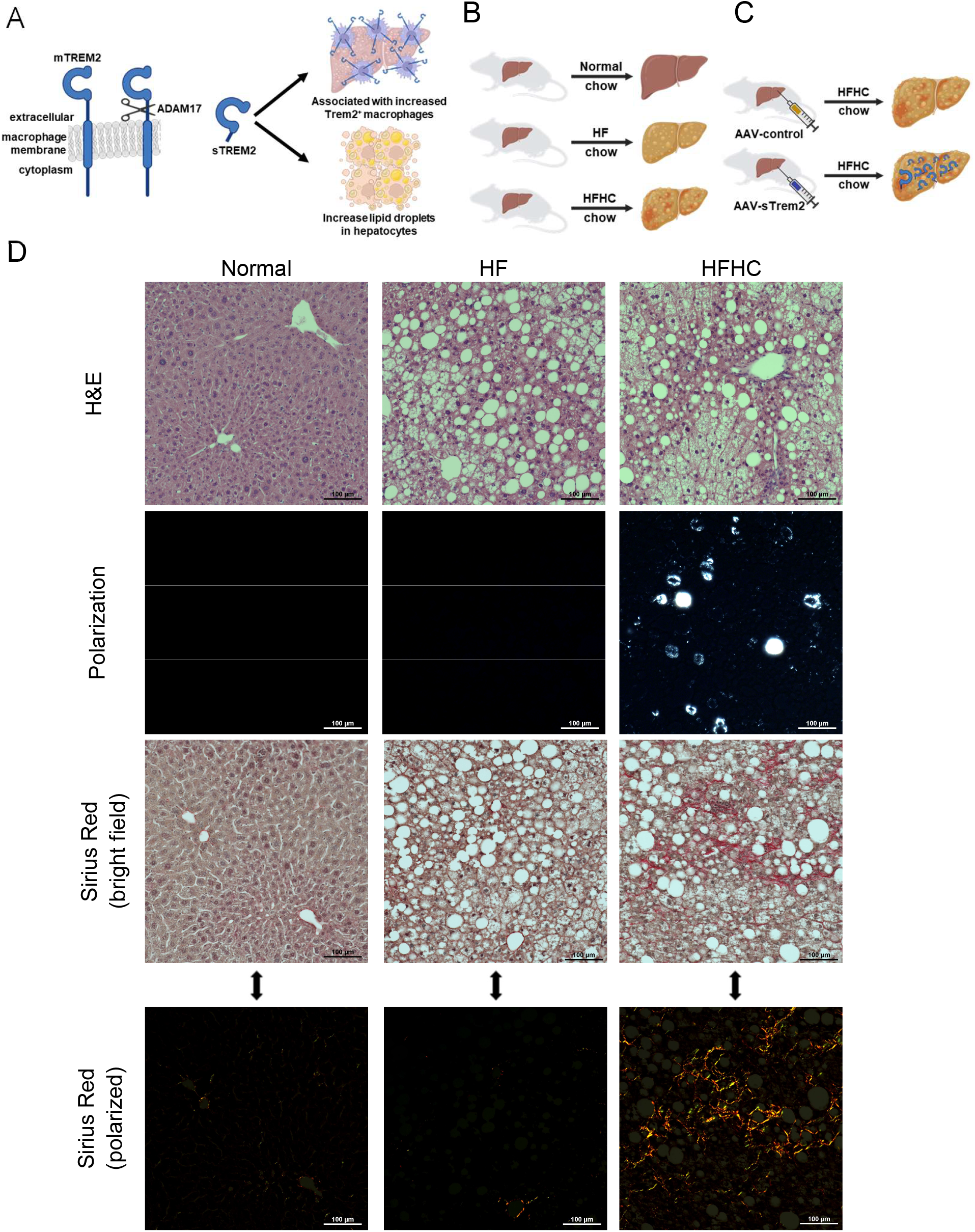
Study design (A-C) and demonstration of the development of fibrosing MASH with cholesterol crystallization in lipid droplets in mice fed a high-fat, high-cholesterol diet (D). **A)** Diagram showing the structure of TREM2 on the membrane of a macrophage. TREM2 is a membrane-bound protein (mTREM2) with an extracellular, lipid-sensing immunoglobulin domain, a membrane bound domain, and an intracellular C-terminus. The extracellular domain can be cleaved by ADAM17 to shed the immunoglobulin domain as soluble TREM2 (sTREM2) which can be detected in plasma. sTREM2 levels are associated with increased Trem2+ macrophages in the liver and increased lipid droplets in hepatocytes **B)** In our first experiment, C57BL/6J mice were exposed to normal, high-fat (HF), or high-fat, high-cholesterol (HFHC) diets, in order to determine if Trem2^+^ macrophages form crown-like structures (CLSs) in the setting of fibrosing MASH with cholesterol crystallization. **C)** In the second experiment, C57BL/6J mice were injected with AAV-control or AAV-sTrem2 (sTrem2 overexpression by hepatocytes) and fed an HFHC diet for 180 days to determine if sTrem2 plays any direct role in the development of fibrosing steatohepatitis. **D)** Histological comparison of C57BL/6J mice assigned to normal, HF, or HFHC diet for 180 days. Liver sections were stained with hematoxylin and eosin (H&E) or Sirius Red. Also, unstained frozen liver sections were viewed under polarized light to identify birefringent cholesterol crystals. Samples were imaged at 200x magnification under brightfield and polarized light. HF diet resulted in severe hepatic steatosis without substantial fibrosis or cholesterol crystallization. HFHC diet resulted in hepatic steatosis with substantial fibrosis (fibrosing MASH), with prominent cholesterol crystallization within lipid droplets.

We aimed to better understand the role of Trem2^+^ macrophages and sTrem2 in the evolution of fibrosing steatohepatitis in an experimental mouse model of MASH (**Figure 1**). We hypothesized that increased sTrem2 protects against cholesterol-induced fibrosing steatohepatitis.

## METHODS

### High-fat (HF) and high-fat, high-cholesterol (HFHC) dietary mouse models of steatotic liver disease

All animal studies were approved by the Animal Care and Use Committee of the Veterans Affairs Puget Sound Health Care System (Seattle, WA, USA). Male, 6-month-old C57BL/6J mice (Jackson Laboratory) were assigned to three different diets for up to 6 months: normal diet (Rodent Chow; PicoLab #5053), high-fat diet (HF; 15% fat, wt/wt; Bio-Serv F6060), or high-fat and high-cholesterol diet (HFHC, 15% fat and 1% dietary cholesterol; Bio-Serv F5947). Cocoa butter, which contains approximately 60% saturated fat, was the source of the extra fat in the diets (**Supplemental Table 1**).

We previously demonstrated that C57BL/6J mice develop fibrosing steatohepatitis after 6 months on the HFHC diet whereas mice on the HF diet develop hepatic steatosis without substantial inflammation or fibrosis [18]. Mice were randomly assigned to be fed the Normal, HF, or HFHC diet for 15 days (n=4-6), 30 days (1 month; n=4-6), 90 days (3 months; n=4-6), or 180 days (6 months; n=4-7). All mice started the diet on the same day (**Figure 1B)**. At the end of each feeding period, mice underwent phlebotomy and were euthanized by cervical dislocation following isoflurane anesthesia. Livers were collected after perfusion.

### TBG-AAV8 vector to induce sTrem2 expression in hepatocytes

An adeno-associated virus serotype 8 (AAV8) vector encoding for the extra-cellular, cleaved portion of Trem2 (i.e., the soluble Trem2 or sTrem2) under the control of the liver-specific human thyroxine binding globulin promoter (TBG-AAV8-sTrem2) was generated (**Supplemental Figure 1**). The identical AAV8 without the sTrem2 sequence (TBG-AAV8-control) was used as a control. The AAV serotype 8 exhibits murine hepatocyte tropism following peripheral vein infusion and is not known to cause disease or immune reactions [26, 27]. We previously used the liver-specific AAV8 in a type 1 diabetes mouse model [28, 29]. The vectors were generated by the CRISPR, Vector and Transgenic Mouse Core (University of Washington Diabetes Research Center, Seattle, WA). Six-month-old, male C57BL/6J mice were treated with the AAV-sTrem2 or AAV-control. The AAVs were injected via the retro-orbital venous sinus at 1 × 10^11^ viral particles in 100 µL/mouse. Two weeks after AAV injections, all mice were fed the HFHC diet. At 15 days, 3 months, and 6 months after starting the HFHC diet (n=10-12 male mice per group), mice were anesthetized by isoflurane followed by phlebotomy, cervical dislocation, and collection of perfused liver by flash freezing in liquid nitrogen (**Figure 1C**). Mouse plasma was isolated for routine blood chemistry analysis. Plasma sTrem2 was quantified using a TREM2 ELISA kit from Raybiotech (Norcross, GA) in 180-day mice.

### Hepatic gene expression studies by RNA sequencing (RNA-seq)

Total hepatic RNA was isolated from frozen mouse livers using RNeasy kits (Qiagen) with on-column DNase digestion. RNA (0.5 ng) was reverse transcribed into full length amplified cDNA. Dual-index, single-read sequencing of pooled libraries was conducted on a HiSeq2500 sequencer (Illumina) with a target depth of 5 million reads per sample. Raw image files were converted to sequenced reads (fastq) with CASAVA base recognition (Illumina). Sequenced reads were aligned to the mouse reference genome GENCODE vM27 using HISAT2 [30], and gene counts were quantified using featureCounts [31]. There was an average of 19.4 million mapped counts per sample. Transcript count data was imported into R and transcripts were filtered for annotated protein-coding genes with at least 10 counts in at least one sample. Transcript normalization and statistical analysis was performed with DESeq2 using the Benjamini-Hochberg false discovery rate (BH-FDR < 0.05) [32]. Relative gene expression was visualized using ComplexHeatmap without clustering.

### Histology of mouse liver tissue

Sections of formalin-fixed, paraffin embedded mouse liver (10 μm) were stained with hematoxylin and eosin (H&E), or Sirius Red. H&E slides were used to determine steatosis scores. Sirius Red was used to visualize fibrosis using both bright field and polarizing microscopy (Nikon NiE). The amount of fibrosis was measured using the polarized images with Image J software. To identify free cholesterol, frozen liver sections (10 μm) were stained with filipin, which interacts with the 3β-hydroxy group of free cholesterol to fluoresce blue, coverslipped with glycerol and then viewed under a DAPI filter of a fluorescent microscope. For visualization of birefringent cholesterol crystals, frozen liver sections (10 μm) were viewed using polarizing microscopy and quantified using Image J software.

### Immunohistochemistry for Trem2+ and Cd68+ macrophages in mouse liver tissue

Mouse livers flash frozen in optimal cutting temperature (OCT) medium were cut to 10 µm sections and fixed with formalin for 10 minutes. Sections were blocked overnight (3% BSA and 0.05% Tween 20 in PBS) and incubated with antibodies for Trem2 (Invitrogen PA5-46978 which recognizes the extracellular domain of and soluble portion of Trem2; 2 µg/mL) or Cd68 (Invitrogen PA5-78996; 175x dilution) for 2 hours followed by washing. Secondary antibodies IgG-AlexaFluor 594 (Invitrogen A11016; 2 µg/mL) or IgG-AlexaFluor 488 (Invitrogen A11008; 1 µg/mL), respectively, were incubated for 1 hour followed by washing. Sections were then coverslipped using Aquamount containing 10 µM Hoechst 33342 as a nucleus stain. All photos were taken by fluorescence microscopy (Nikon NiE microscope or Nikon A1R confocal microscope) at 200x original magnification. The percentage of positive staining was calculated using Image J analysis software.

### Statistical analysis

Students t-test was used to compare three diet groups (normal versus HF; normal versus HFHC; and HF versus HFHC) or to compare AAV-sTrem2 versus AAV-Control mice with respect to mean values of blood chemistry, histology, and other results. A p-value <0.05 was considered statistically significant.

## RESULTS

### HF diet causes steatosis but HFHC diet causes fibrosing steatohepatitis with cholesterol crystallization

Mice on a HF diet gained weight and developed severe macrovesicular hepatic steatosis with elevated liver transaminases without substantial fibrosis or any hepatic cholesterol crystallization within lipid droplets (**Figure 1D and Table 1**). HF- and HFHC-fed mice gained a similar amount of weight associated with severe macrovesicular hepatic steatosis and elevated liver transaminases. However, compared to HF-fed mice, HFHC-fed mice had greater liver weight and liver/mouse weight ratio (**Table 1**). After 90 days on a HFHC diet, there was no evidence of hepatic fibrosis or cholesterol crystallization (**Supplemental Figure 2**). After 180 days, the HFHC-fed mice developed substantial hepatic fibrosis (i.e., fibrosing steatohepatitis) as well as prominent birefringent crystals within lipid droplets consistent with cholesterol-containing crystals (**Figure 1D, Table 1**).

**Table 1.**
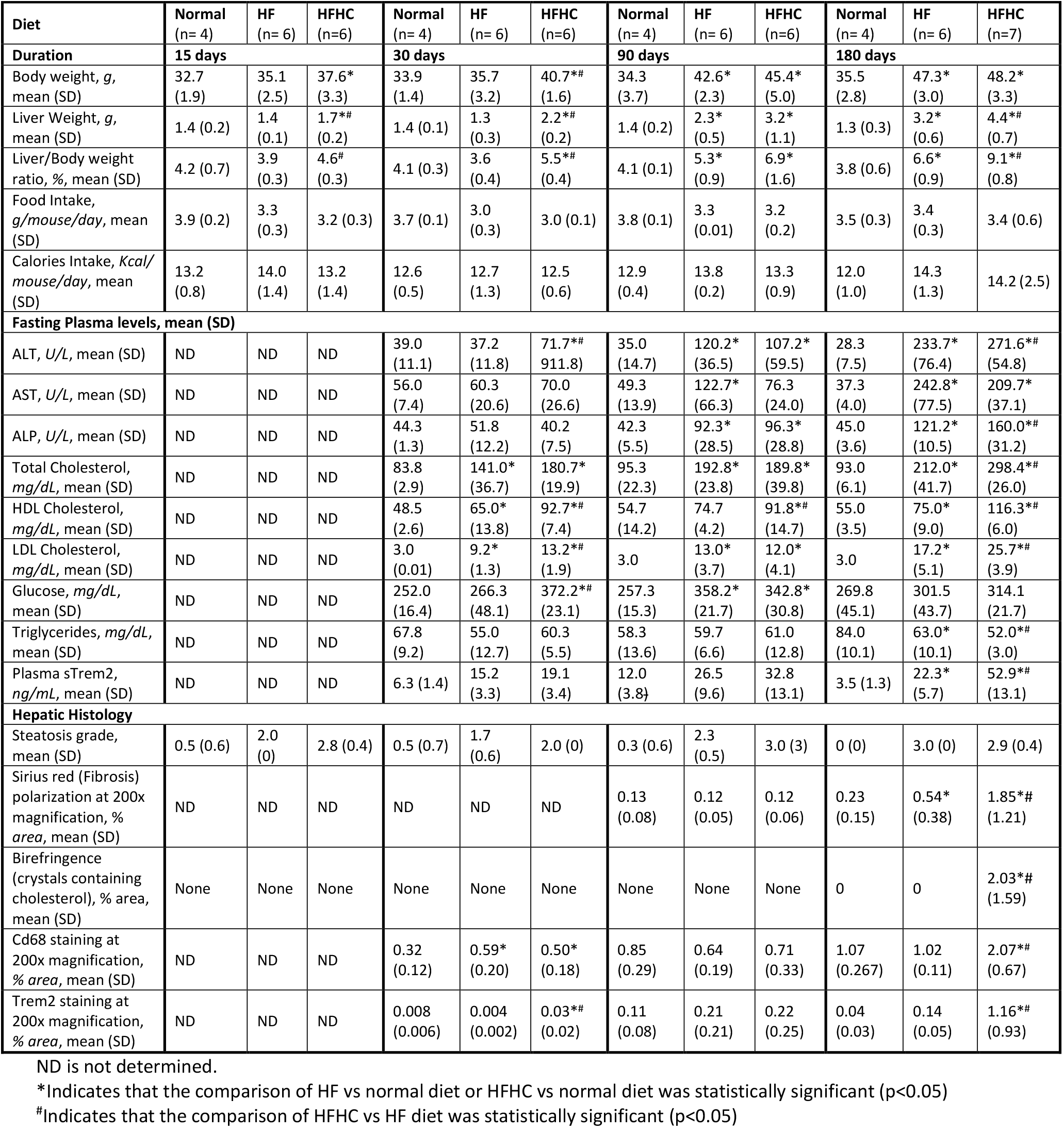
Comparison of the C57BL/6J mice assigned to normal diet, HF diet or HFHC diet for up to 6 months and sacrificed at 15, 30, 90 and 180 days from diet initiation for physical attributes, blood chemistry, and histology.

### HFHC diet causes the development of crown-like structures (CLS) consisting of Trem2+ macrophages

In contrast to normal- and HF-fed mice, HFHC-fed mice had increased macrophage staining for Trem2 and Cd68 at 180 days into the assigned diets (**Table 1**). Visually, Trem2 and Cd68 fluorescence formed circular CLSs around necrotic, steatotic hepatocytes and their remnant lipid droplets (**Figure 2**). These circular structures of macrophages were absent from the normal- and HF-fed mice. In mice fed a normal or HF-diet, Cd68 staining was scattered with no observable structural features, and Trem2 staining was minimal to not-detectable in HF- and normal-fed mice, respectively. In HFHC-fed mice at 180 days, co-staining of Trem2 and the nucleus with DAPI showed the highly consistent formation of CLSs by Trem2^+^ macrophages (**Figure 3A**). Trem2^+^ (**Figure 3B**) and Cd68^+^ (**Figure 3C**) macrophages surround the remnant lipid droplets of necrotic hepatocytes forming CLCs around the birefringent, crystalline material that is observed under polarized light. Co-staining with filipin (which stains free cholesterol) demonstrated that the birefringent material contained free cholesterol crystals (**Figure 3D**).

**Figure 2.**
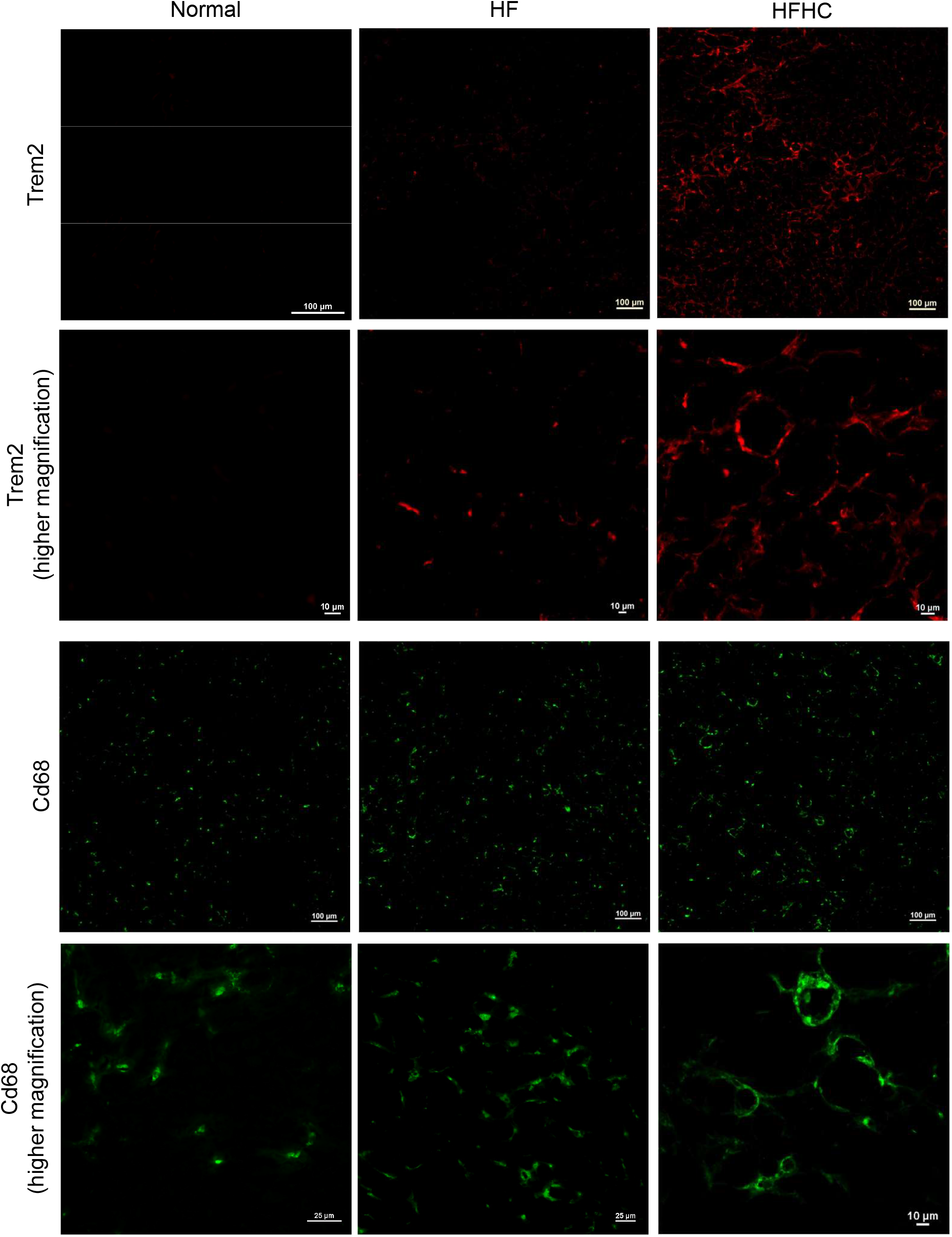
Immunohistochemistry for Trem2 and Cd68 comparing liver sections of mice fed normal, high-fat (HF), and high-fat, high-cholesterol (HFHC) diet for 180 days. Comparison of Trem2 (red fluorescence) and Cd68 (green fluorescence) in mice fed a normal, HF, and HFHC diet for 180 days. The scale bar at the bottom right of each image is 10-100 µm. Images at higher magnification of liver sections in HFHC-fed mice show Trem2+ and CD68+ macrophages forming prominent circular crown-like structures (CLSs), which are not seen in normal-or HF-fed mice.

**Figure 3.**
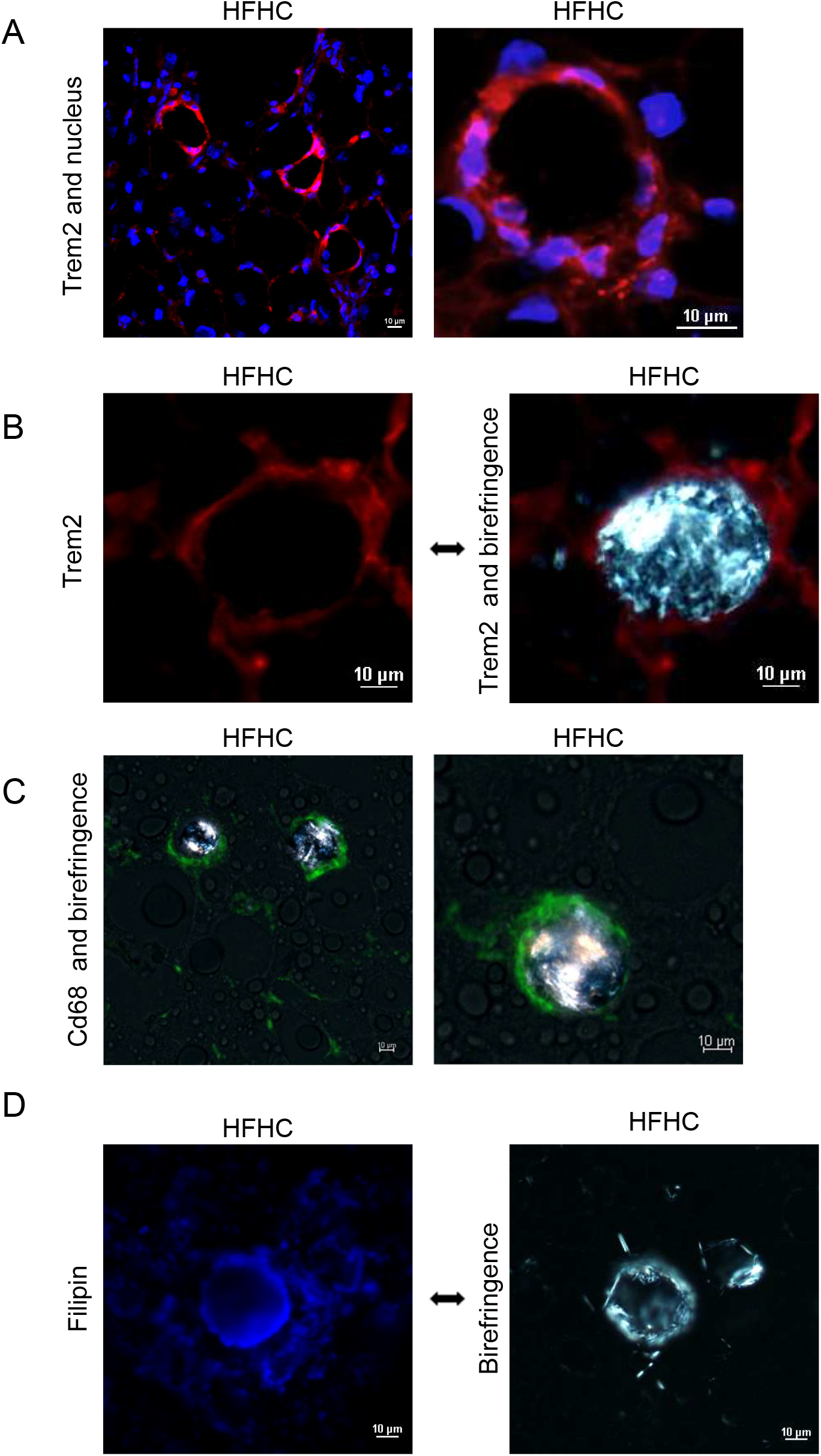
Crown-like structures of Trem2+ macrophages surround necrotic, steatotic hepatocytes which contain cholesterol crystals in their lipid droplets, in mice fed a HFHC diet for 180 days. **A)** Co-fluorescence of Trem2 (red) and the nucleus (blue). Scale bars are 10 µm. **B)** Trem2 fluorescence (red) compared to the same Trem2 fluorescence with birefringence (light polarization) overlay to demonstrate the circular crown-like structures formed by Trem2 surrounding the birefringent cholesterol crystals. Scale bars are 10 µm. **C)** Image overlays of Cd68 fluorescence (green) and birefringence to demonstrate the circular crown-like structures formed by Cd68 surrounding the birefringent cholesterol crystals. Scale bars are 10 µm. **D)** Free cholesterol fluorescence (filipin) compared to birefringent cholesterol crystals using the same image location. Scale bars are 10 µm.

### Plasma sTrem2 levels track liver disease progression from normal to steatosis to fibrosing steatohepatitis (MASH)

Plasma sTrem2 levels tracked the progression of steatohepatitis both over time and across models (**Table 1**). Plasma sTrem2 levels increased in mice on HFHC diet from 19.1 ng/mL after 30 days on HFHC diet to 32.8 ng/mL after 90 days (when severe steatosis had developed but no fibrosis or cholesterol crystallization) to 52.9 ng/mL after 180 days on a HFHC diet (when fibrosing steatohepatitis had developed together with cholesterol crystallization). There was no difference in sTrem2 levels between HFHC and HF-fed mice at 90 days (32.8 vs. 25.5) when both groups developed steatosis without fibrosis or cholesterol crystals, but at 180 days, plasma sTrem2 concentrations were much higher in the HFHC-fed mice that developed fibrosing steatohepatitis than in the HF-fed mice that did not (52.9 ng/mL vs. 22.3 ng/mL). Overall, by day 180, plasma sTrem2 levels were 6-fold higher in the HF-fed mice (22.3 ± 5.7 ng/mL) than in the normal-diet mice (3.5 ± 1.3 ng/mL), but they were 15-fold higher in the HFHC-fed mice (52.9 ± 13.1 ng/mL).

### HFHC diet increases hepatic RNA expression of genes associated with inflammation, including Trem2

Compared to normal-fed mice HF- and HFHC-fed mice had increased hepatic RNA expression of inflammation and myeloid genes, as well as fibrosis related genes (**Figure 4 and Supplemental Table 2**). At 180 days, compared to HF-fed mice, HFHC-fed mice had increased RNA expression for genes related to inflammation and myeloid cell activation (*Trem2, Cd68, Csf1* and *Tyrobp*), liver fibrosis (*Lgals3* and *Mmp12*), and sterol export (*Abcg5* and *Abcg8*). Compared to normal diet, macrophage-expressed genes *Aif1* and *Adgre1* had increased RNA expression at 90 and 180 days, indicating that HFHC mice compared to HF mice may have increased *Trem2*+ expression independent of increased macrophages. Compared to normal- and HF-fed mice, HFHC-fed mice had decreased expression of genes involved in cholesterol synthesis, biotransformation, and regulation (*Pcsk9, Hmgcr, Hmgcs1, Sqle, Insig*, and *Cyp8b1*), indicative of cholesterol accumulation. Compared to normal-fed mice, HF- and HFHC-fed mice had increased RNA expression of the lipid and cholesterol storage related genes *Plin3, Plin4, Soat1*, and *Scap* at 180 days. Taken together, compared to normal-fed and HF-fed mice, HFHC-fed mice had increased expression of pro-inflammatory, pro-fibrotic, and sterol export genes, as well as decreased sterol synthesis genes.

**Figure 4.**
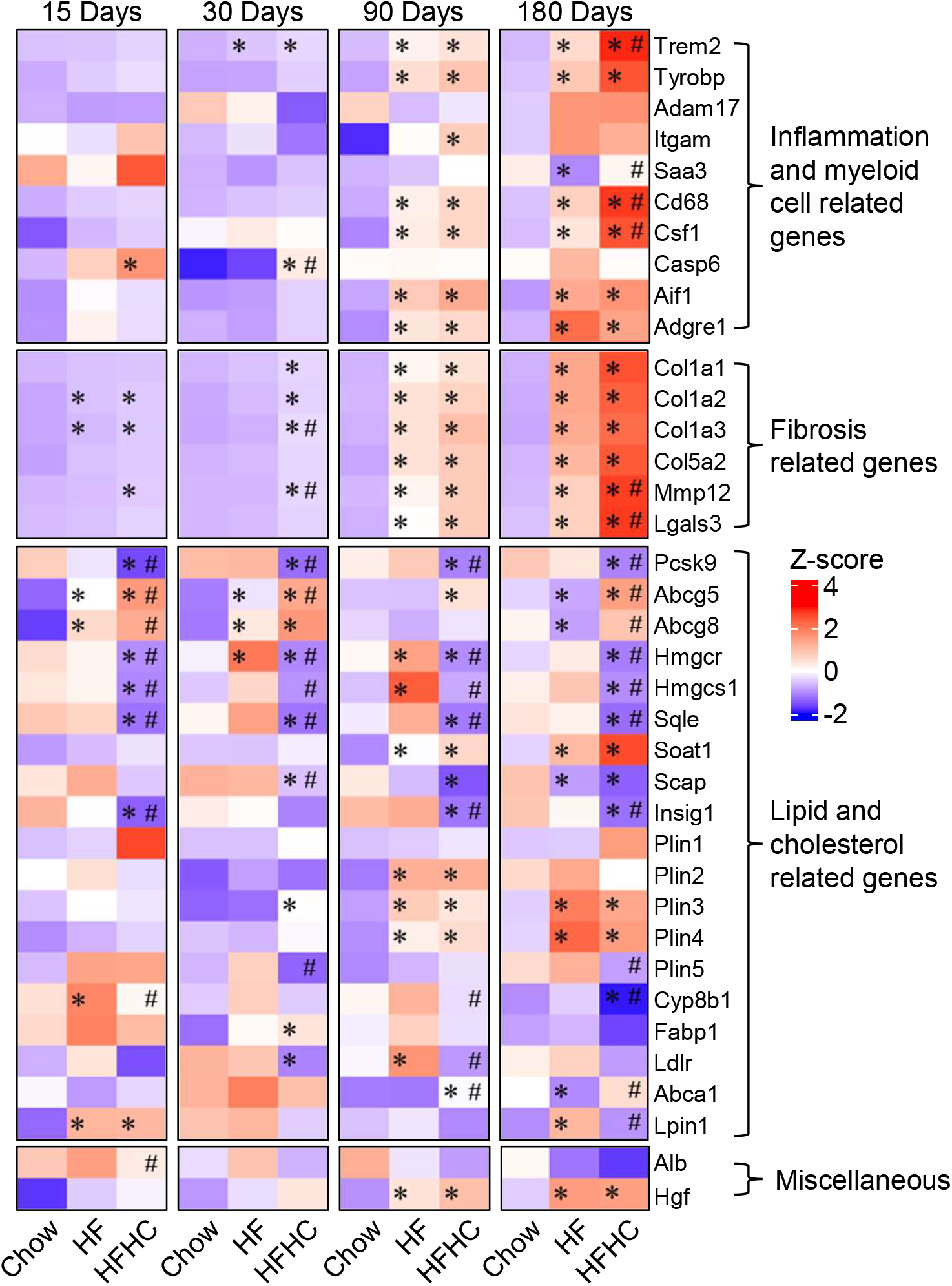
Relative hepatic RNA expression of inflammation, fibrosis, and lipid genes comparing mice on normal, HF, and HFHC diets. Heatmap showing the relative hepatic RNA expression of selected inflammation, fibrosis, and lipid genes in mice fed normal, HF, or HFHC diet for 15, 30, 90, and 180 days. A “*” indicates q<0.05 compared to Chow within an age group. A “#” indicates q<0.05 compared to HF within an age group. The normalized counts are provided in **Supplemental Table 2**.

### AAV-sTrem2 mice exhibit higher plasma sTrem2 levels than AAV-control mice but no increase in steatosis or fibrosis

Mice injected with AAV-sTrem2 had more than two times higher plasma sTrem2 levels than mice injected with AAV-control (138.7 ng/mL vs. 51.5 ng/mL), indicating that the AAV transduction was successful and resulted in higher production of sTrem2 by transduced hepatocytes (**Table 2**). However, the AAV-sTrem2 and AAV-control mice had similar body weight, liver weight, liver/body weight ratio, plasma levels of transaminases, and lipids, hepatic steatosis, cholesterol crystals and fibrosis after 180 days (**Table 2** and **Figures 5A, 5B, and 5C**) and at earlier time points (15 days and 90 days; **Supplemental Table 3**). It was noted that both mice injected with AAV-sTrem2 and those injected with AAV-control had similarly low levels of fibrosis, which were lower than in our experiments shown in **Table 2** in which no AAV was injected. Compared to AAV-control mice, AAV-sTrem2 had increased Trem2 staining but similar Cd68 staining at 180 days (**Table 2** and **Figures 5D and 5E**). CLSs were prominent by both Trem2 and Cd68 fluorescence in both AAV-sTrem2 and AAV-control mice. Taken together, the hepatocyte-targeted AAV overexpression of sTrem2 did not alter the progression of steatohepatitis in HFHC-fed mice.

**Table 2.** Comparison of C57BL/6J mice injected with AAV-sTrem2 versus AAV-control and then assigned to HFHC diet for 180 days for physical attributes, blood chemistry, and histology.

|  | 180 days |  |
| --- | --- | --- |
|  | AAV-control<br>(n= 12) | AAV-sTrem2<br>(n= 12) |
| Body weight, *g*, mean (SD) | 46.7 (4.4) | 47.1 (2.0) |
| Liver Weight, *g*, mean (SD) | 3.7 (0.8) | 3.6 (0.6) |
| Liver/Body weight ratio, %, mean (SD) | 7.8 (1.1) | 7.5 (1.1) |
| Food Intake, *g/mouse/day*, mean (SD) | 3.1 (0.3) | 3.3 (0.2) |
| Calories Intake, *Kcal/mouse/day*, mean (SD) | 12.9 (1.2) | 13.7 (0.8) |
| ALT, *U/L*, mean (SD) | 279.8 (86.7) | 335.7 (201.7) |
| AST, *U/L*, mean (SD) | 212.9 (63.2) | 231.9 (125.9) |
| ALP, *U/L*, mean (SD) | 181.7 (34.2) | 150.5 (39.8) |
| Total Cholesterol, *mg/dL*, mean (SD) | 244.2 (25.8) | 210.7 (51.2) |
| HDL Cholesterol, *mg/dL*, mean (SD) | 116.3 (8.1) | 106.3 (21.7) |
| LDL Cholesterol, *mg/dL*, mean (SD) | 19.9 (4.2) | 18.6 (5.3) |
| Glucose, *mg/dL*, mean (SD) | 297.7 (53.9) | 256.3 (42.4) |
| Triglycerides, *mg/dL*, mean (SD) | 57.0 (5.4) | 52.3 (12.0) |
| Plasma sTrem2, *ng/mL*, mean (SD) | 51.5 (12.2) | 138.7* (10.9) |
| Steatosis grade, mean (SD) | 2.9 (0.3) | 3.0 (0.0) |
| Sirius red (Fibrosis) polarization at 200x magnification, % *area*, mean (SD) | 0.15 (0.11) | 0.14 (0.12) |
| Birefringence (crystals containing cholesterol), % *area*, mean (SD) | 0.538 (0.742) | 0.399 (0.522) |
| Cd68 staining at 200x magnification, % *area*, mean (SD) | 0.628 (0.480) | 0.997 (0.325) |
| Trem2 staining at 200x magnification, % *area*, mean (SD) | 0.119 (0.075) | 0.701** (0.402) |

\*Indicates that the comparison of AAV-Trem2 vs AAV-Control was statistically significant (\*p<0.05; \*\*p<0.01)

**Figure 5.**
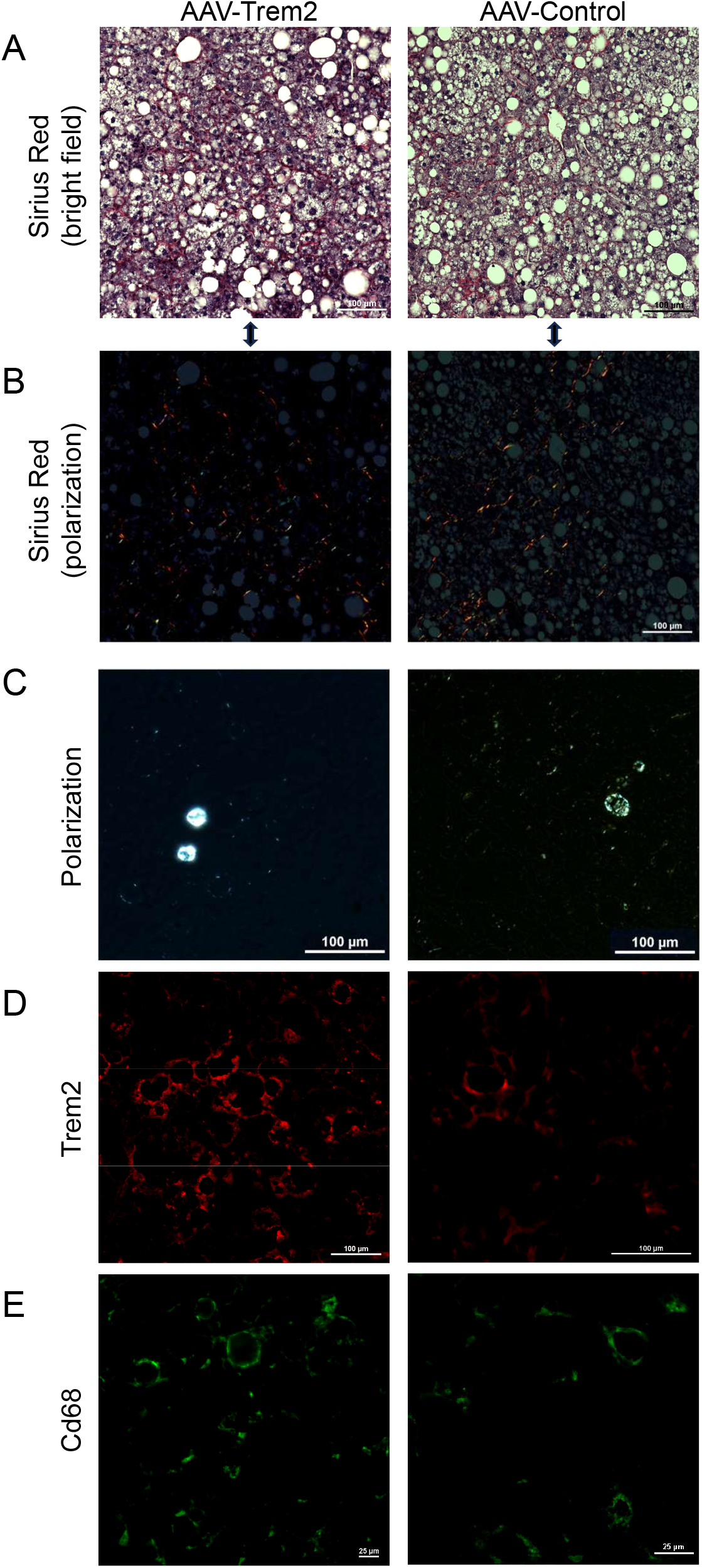
Histological comparison of mice injected with AAV-sTrem2 versus AAV-control and were then fed a HFHC diet for 180 days. Liver from mice injected with AAV-sTrem2 and AAV-control were fed an HFHC diet for 180 days. **A)** Histology with Sirius Red staining under bright field and **B)** Sirius Red staining under polarization. Percentage area values are provided in **Table 2. C)** Histology samples imaged under polarized light demonstrating similar levels of birefringence (crystals containing cholesterol). **D)** Trem2 fluorescence (red) was greater in AAV-Trem2 mice compared to AAV-control mice. **E)** Cd68 fluorescence (green) was similar in AAV-Trem2 and AAV-control mice.

## DISCUSSION

In these animal experiments, we demonstrated that during the development of fibrosing MASH, Trem2^+^ macrophages selectively formed conspicuous CLSs surrounding and processing steatotic, necrotic hepatocytes and their remnant lipid droplets, which contained cholesterol crystals (**Figure 6A**). This suggests that Trem2^+^ macrophages play a vital role in fibrosing MASH through the development of CLSs that process the lipids in remnant lipid droplets, including cholesterol crystals. Furthermore, plasma sTrem2 levels tracked the progression of disease over time and across dietary models, increasing progressively from normal liver to steatosis (6-fold increase) to fibrosing steatohepatitis/MASH (15-fold increase). However, overexpression and secretion of sTrem in hepatocytes caused by AAV did not modify the development of steatosis or fibrosis in MASH (**Figure 6B**). This suggests that sTrem2, which is normally cleaved from membrane-bound Trem2 and can be measured in plasma as a promising biomarker of MASH, does not itself have a direct impact on the development of fibrosing steatohepatitis.

**Figure 6.**
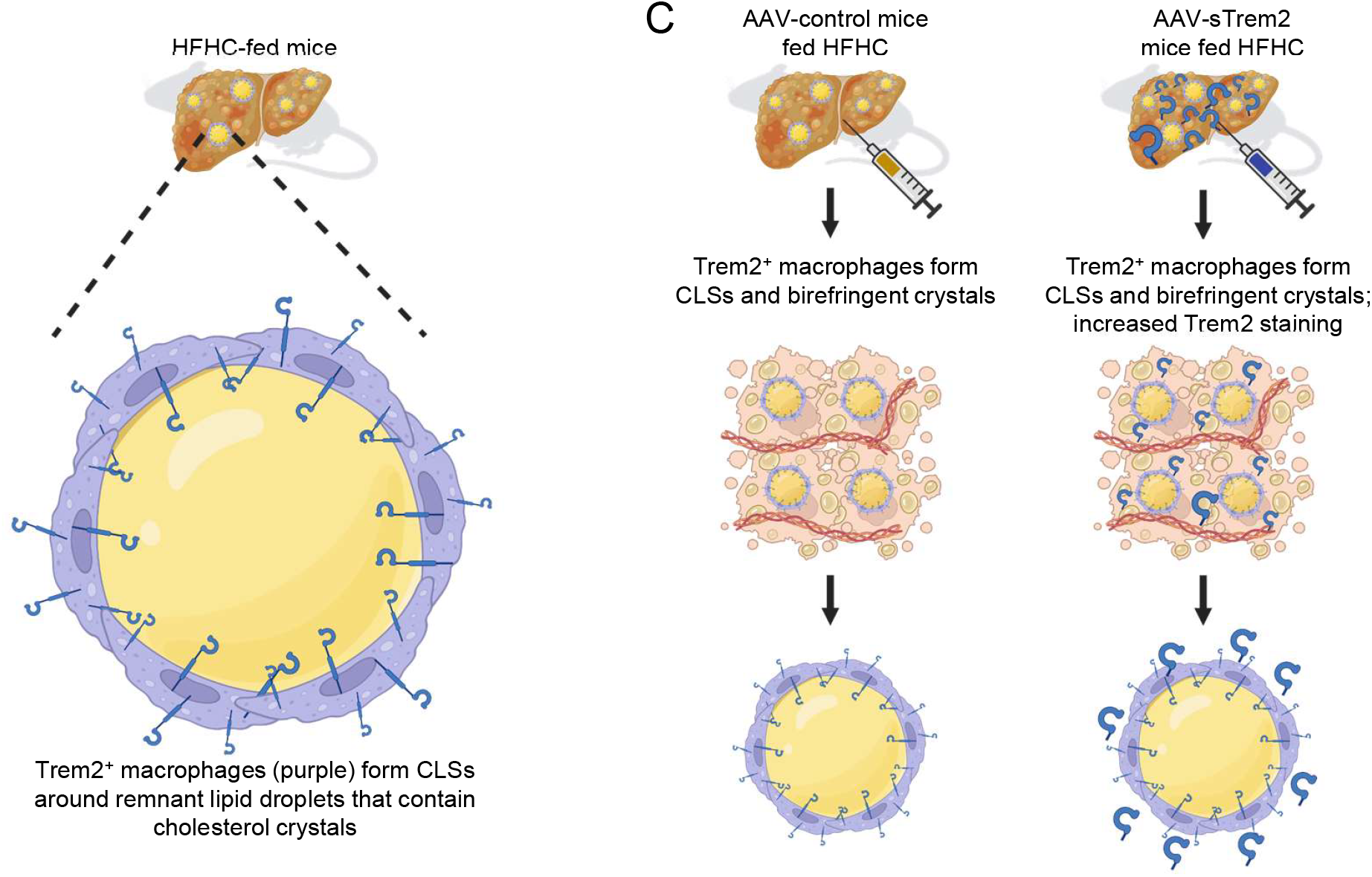
Schematic representation of the results. **A)** Trem2+ macrophages forming crown-like structures surrounding lipid droplets that contain cholesterol crystals in HFHC-fed mice. **B)** Illustration that sTrem2 overexpression in AAV-sTrem2 injected mice did not affect the development of steatosis or fibrosis in MASH compared to AAV-control injected mice.

Trem2^+^ macrophages accumulate around damaged steatotic hepatocytes and their remnant lipid droplets to form CLSs— likely in response to lipid molecules in the remnant lipid droplets since Trem2 is known to be a lipid-sensing receptor. We demonstrated that cholesterol crystals are frequently found at the center of these CLSs, as well as the other known lipid components of lipid droplets, such as triglycerides, cholesterol esters, and phospholipids. TREM2^+^ macrophages are responsible for clearing cholesterol debris and other lipids in necrotic, steatotic hepatocytes to protect the liver from further lipotoxic cellular damage [18]. They have a unique transcriptional signature—conserved across tissue types—that is best suited to phagocytose and process lipids [11, 33]. Compared to wild-type mice, Kupffer cells and infiltrating macrophages of Trem2-deficient mice can maintain uptake of triglyceride-rich lipoproteins; however, Trem2-deficient bone marrow-derived macrophages had decreased capacity to phagocytose apoptotic hepatocytes, indicating that efferocytosis in hepatic steatosis is Trem2-dependent [11]. In our fibrosing steatohepatitis MASH mouse model induced by HFHC diet, the Trem2^+^ CLSs indicate sites of efferocytosis of damaged, steatotic hepatocytes and subsequent processing of their lipid content, including cholesterol crystals.

A preponderance of evidence suggests that TREM2^+^ macrophages protect against steatosis-induced liver injury

TREM2^+^ macrophages remove nonfunctional hepatocytes and lipids, inhibit inflammation and degrade collagen, indicating that TREM2^+^ macrophages may be associated with MASH regression [14, 15, 34]. Furthermore, TREM2 deficiency or ablation is extensively associated with defective lipid metabolism, increased inflammation, and exacerbated steatohepatitis [11, 14, 15]. However, TREM2^+^ macrophages were also found to co-express the peptide membrane spanning 4-domains A7 (MS4A7) that drives steatohepatitis through nucleotide-binding oligomerization domain-like receptor family pyrin domain containing 3 (NLRP3) inflammasome activation [35]. The basis for TREM2^+^ macrophages capable of both pro- and anti-inflammatory signaling is not known. It is also possible that TREM2^+^ macrophages may not be a homogeneous class of macrophages and may contain subsets mediating different biologic functions. Reconciling the lineages and subtypes of TREM2^+^ macrophage may be key to regulating MASH regression and ameliorating steatosis-induced fibrosis.

The hepatocyte-targeted overexpression of sTrem2 induced by an AAV8 vector did not appear to mediate significant effects on the liver of HFHC-induced MASH mice (**Table 2** and **Figure 5A**). In our hepatocyte-targeted, overexpression model of sTrem2, both AAV-control and AAV-sTrem2 mice developed fibrosing steatohepatitis and CLSs with birefringent cholesterol crystals equally. However, the percentage area of fibrosis in the AAV-control mice (**Table 2**) was less than the HFHC mice (**Table 1**). This may indicate that AAV8 alters hepatic immune response to steatosis and demonstrates the variability within fibrosing steatohepatitis models. An increase in plasma sTrem2 level did not alter the kinetics of developing fibrosing steatohepatitis in the liver. Others have found that—in Trem2-deficient mice—the addition of recombinant sTrem2 did not protect mice from MASH [15]. Indeed, our model, which has endogenous sTrem2 production, provides added evidence that sTrem2 does not directly impact MASH development. An increase in plasma sTrem2 is found to be associated with increased Trem2^+^ macrophages aggregating around pro-inflammatory, fibrotic regions of the liver— an observation believed to protect against the development of MASH [14]. It is unclear if sTrem2 is directly involved in the lineage determination of Trem2^+^ macrophages or if the increased plasma sTrem2 levels simply reflect higher numbers of Trem2+ macrophages which result in higher levels of plasma sTrem2 after cleavage and release.

We observed increased Trem2^+^ fluorescence and Trem2^+^ CLSs in AAV-sTrem2 mice, but both AAV-sTrem2 and AAV-Control mice had similar levels of fibrosis (Table 2) that were below the fibrotic levels of mice with no AAV injection. It is unclear if artificially increasing sTrem2 would protect mice from fibrosis compared to control mice if the diet-induced steatosis was extended past 180 days. It is possible that the sTrem2 function is naturally saturated in diet-induced MASH models without artificially increasing sTrem2, preventing an agonistic, protective effect by the AAV-sTrem2. Furthermore, in the human brain of Alzheimer’s disease patients, there is a heterogenous pool of sTrem2 arising from alternatively spliced isoforms of the *TREM2* transcript [36]. Alternative sTrem2 isoforms in a MASH liver have not been reported. More research is needed to understand the interplay of mTrem2 and sTrem2 in MASH.

In conclusion, mice fed HFHC diet developed fibrosing steatohepatitis (MASH) in association with a 15-fold increase in plasma sTrem2 levels corresponding to Trem2^+^ macrophages forming prominent CLSs around steatotic, necrotic hepatocytes that containing cholesterol crystals in their lipid droplets. Elevated plasma sTrem2 may serve as a biomarker for CLSs and fibrosing steatohepatitis, but engineered overexpression of sTrem2 did not ameliorate hepatic steatosis or fibrosis and is therefore unlikely to exert beneficial effects on the liver in MASH.

## List of abbreviations

(AAV): Adeno-associated viral vector
AST: aspartate aminotransferase,
ALT: alanine aminotransferase,
CLSs: crown-like structures,
HF diet: high-fat diet,
HFHC diet: high-fat, high-cholesterol diet, metabolic dysfunction-associated steatotic liver disease,
MASLD: metabolic dysfunction-associated steatohepatitis,
MASH: triggering receptor expressed on myeloid cells 2,
TREM2: soluble TREM2, sTREM2.

