## Supplemental appendix for "Soluble TREM2 is a biomarker but not a mediator of fibrosing steatohepatitis"

**Supplemental table 1.** Composition of the diets used in the experiments.

| | Normal | HF | HFHC |
|---|---|---|---|
| **Protein, g/ 100 g** | 21.0 | 14.4 | 14.4 |
| **Carbohydrate, g/ 100g** | 65.2 | 57.1 | 56.1 |
| **Fat, g/ 100 g** | 5.0 | 15.2 | 15.2 |
| **Cholesterol, g/100g** | < 0.02 | < 0.02 | 1.0 |
| **Fatty Acids, g/100g** | | | |
| C18:2 | 2.68 | 1.4 | 1.4 |
| C18:3 | 0.06 | 0.2 | 0.1 |
| Total Saturated | 0.77 | 8.1 | 8.1 |
| Total Monounsaturated | 1.0 | 4.8 | 4.8 |
| Total Polyunsaturated | 2.73 | 1.6 | 1.6 |
| **Protein, kcal/g** | 0.7 | 0.6 | 0.6 |
| **Carbohydrate, kcal/g** | 2.6 | 2.3 | 2.2 |
| **Fat, Kcal/g** | 0.5 | 1.4 | 1.4 |
| **Total kcal/g** | 3.8 | 4.2 | 4.2 |

The normal diet is Picolab Rodent diet #5053. The high-fat (HF) diet is Bio-Serv HF Diet #F6060, and the high-fat, high-cholesterol diet is Bio-Serv HFHC Diet #F5947.

**Supplemental table 2.** Hepatic genes expression (normalized expression) of C57BL/6J mice fed a normal, HF diet, or HFHC diet.

| Duration | 15 days | | | 30 days | | | 90 days | | | 180 days | | |
|---|---|---|---|---|---|---|---|---|---|---|---|---|
| Diet | Normal n=2 | HF n=3 | HFHC n=4 | Normal n=3 | HF n=3 | HFHC n=3 | Normal n=2 | HF n=3 | HFHC n=4 | Normal n=3 | HF n=4 | HFHC n=3 |
| **Inflammation and Myeloid Cell Related Genes** | | | | | | | | | | | | |
| Trem2 | 14.1 | 16.0 | 26.6 | 1.1 | 14.2* | 25.6* | 9.0 | 68.5* | 87.8* | 10.4 | 133.6* | 367.7*# |
| Tyrobp | 66.3 | 96.9 | 112.4 | 53.1 | 54.9 | 90.4 | 54.0 | 186.6* | 224.6* | 112.8 | 301.9* | 516.8* |
| Adam17 | 355.1 | 349.7 | 348.4 | 366.2 | 347.5 | 299.4 | 368.4 | 330.7 | 340.1 | 457.6 | 539.1 | 537.2 |
| Itgam | 10.2 | 8.9 | 14.8 | | | | 1.2 | 9.6 | 13.0* | 10.1 | 22.9 | 20.4 |
| Saa3 | 105.6 | 66.8 | 144.7 | 35.5 | 28.1 | 39.9 | 36.8 | 41.4 | 57.6 | 89.7 | 33.9* | 84.3# |
| Cd68 | 95.0 | 139.7 | 153.9 | 92.9 | 114.3 | 129.0 | 84.7 | 243.8* | 304.2* | 159.3 | 442.9* | 875.3*# |
| Csf1 | 135.3 | 200.2 | 214.1* | 217.2 | 246.0 | 227.4 | 153.8 | 250.2* | 272.5* | 258.2 | 352.0* | 550.1*# |
| Casp6 | 351.0 | 487.7 | 575.0* | 204.1 | 228.8 | 403.0*# | 387.4 | 391.1 | 384.6 | 529.7 | 664.2 | 521.1 |
| Aif1 | 25.9637 | 51.24304 | 44.57387 | 27.43601 | 29.09736 | 41.55146 | 31.85091 | 75.96788 | 88.50553 | 28.10399 | 89.039 | 98.05877 |
| Adgre1 | 160.829 | 452.9999 | 324.7069 | 228.8113 | 190.5817 | 299.2064 | 146.9822 | 503.5825 | 556.8159 | 233.1721 | 962.7277 | 764.1041 |
| **Fibrosis Related Genes** | | | | | | | | | | | | |
| Col1a1 | 152.2 | 211.8 | 233.6 | 126.3 | 190.3 | 294.1* | 114.6 | 629.6* | 815.9* | 143.3 | 1884.7* | 2841.3* |
| Col1a2 | 152.8 | 285.2* | 324.7* | 148.9 | 238.2 | 330.6* | 214.9 | 786.8* | 895.0* | 230.0 | 1725.1* | 2395.2* |
| Col3a1 | 296.5 | 463.2 | 620.3* | 274.5 | 356.1 | 703.1*# | 364.5 | 1499.3* | 2020.0* | 377.8 | 3188.4* | 4359.0* |
| Col5a2 | 26.5 | 47.7 | 53.6 | 35.3 | 39.1 | 56.1 | 27.4 | 116.8* | 140.0* | 50.8 | 222.5* | 349.3* |
| Mmp12 | 6.2 | 21.0 | 51.0* | 1.2 | 6.2 | 70.3*# | 3.3 | 211.0* | 362.6* | 12.3 | 482.7* | 1153.2*# |
| Lgals3 | 62.4 | 71.1 | 110.3 | 47.1 | 55.6 | 100.8 | 36.4 | 198.4* | 379.3* | 95.0 | 497.2* | 1126.4*# |
| **Lipid and Cholesterol Related Genes** | | | | | | | | | | | | |
| Pcsk9 | 1639.6 | 1052.9 | 263.4*# | 1636.1 | 1659.4 | 400.1*# | 1244.1 | 1550.1 | 503.3*# | 2133.5 | 1773.6 | 694.2*# |
| Abcg5 | 958.9 | 2223.1* | 3700.2*# | 1051.2 | 1799.6* | 3156.5*# | 1600.9 | 1576.2 | 2461.5* | 2345.8 | 1895.0* | 4511.8*# |
| Abcg8 | 1475.6 | 3629.1* | 4288.8# | 1767.7 | 3084.7* | 4140.3* | 2505.1 | 2211.6 | 2602.8 | 4077.5 | 2880.0* | 4904.7# |
| Hmgcr | 2668.1 | 2252.1 | 1018.2*# | 1752.7 | 3837.2* | 871.9*# | 2014.2 | 3327.2* | 927.2*# | 2146.4 | 3037.6* | 1100.3*# |
| Hmgcs1 | 4187.7 | 3739.5 | 1253.8*# | 2202.5 | 4259.8 | 1350.6# | 2128.1 | 7581.7* | 1730.9# | 4932.8 | 6576.3 | 1734.1*# |
| Sqle | 2146.0 | 1949.1 | 139.5*# | 1327.7 | 2451.6 | 145.4*# | 1050.6 | 2310.8 | 192.6*# | 2158.8 | 1894.6 | 117.9*# |
| Soat1 | 50.0 | 70.3 | 92.9 | 69.1 | 68.6 | 90.9 | 36.2 | 101.1* | 142.4* | 106.5 | 237.5* | 368.5* |
| Scap | 3035.3 | 3592.8 | 2444.8 | 3213.7 | 3157.4 | 2193.7*# | 2758.5 | 2194.6 | 1659.7* | 4254.4 | 2779.4* | 2321.7* |
| Insig1 | 6725.2 | 4954.7 | 2790.0*# | 4859.4 | 4535.0 | 2939.6 | 5991.1 | 6349.5 | 2826.9*# | 7920.7 | 6428.9 | 3744.5*# |
| Plin1 | 0.0 | 0.6 | 13.2 | | | | | | | | | |
| Plin2 | 16834.8 | 18388.5 | 15817.1 | 11099.2 | 12858.2 | 11692.6 | 12125.8 | 19180.0* | 19133.3* | 23623.7 | 26536.4 | 20999.0 |
| Plin3 | 901.1 | 1028.4 | 967.6 | 627.8 | 653.7 | 933.8* | 758.9 | 1134.9* | 1041.0* | 1169.0 | 1912.5* | 1705.4* |
| Plin4 | 113.7 | 332.5 | 516.3* | 399.6 | 324.8 | 662.4 | 120.8 | 864.0* | 1037.3* | 650.2 | 2825.2* | 2201.2* |
| Plin5 | 1043.7 | 1499.6 | 1481.2 | 934.7 | 1213.3 | 787.1# | 870.2 | 950.0 | 1025.4 | 1642.8 | 1824.5 | 1228.9# |
| Cyp8b1 | 9173.6 | 13162.9* | 8086.8# | 5827.4 | 9011.1 | 5844.8 | 7551.6 | 10338.8 | 6323.7# | 6166.6 | 8225.5 | 2716.1*# |

| | | | | | | | | | | | | |
|---|---|---|---|---|---|---|---|---|---|---|---|---|
| Fabp1 | 42881.4 | 54051.0 | 46641.5 | 24733.9 | 34968.0 | 37390.1* | 33710.7 | 40610.3 | 32731.4 | 39055.9 | 40876.2 | 30091.7 |
| Ldlr | 4359.7 | 5300.0 | 3614.8 | 5368.5 | 5158.0 | 3635.7* | 4490.5 | 5843.2* | 3831.0# | 6488.2 | 6969.0 | 5232.7 |
| Abca1 | 4148.4 | 3442.0 | 3883.5 | 4749.4 | 5329.5 | 4595.0 | 2959.4 | 2940.0 | 3835.2*# | 5324.2 | 4179.8* | 5833.0# |
| Lpin1 | 1267.0 | 4935.4* | 4832.9* | 4152.6 | 4432.0 | 2358.5 | 2240.8 | 2655.0 | 1573.8 | 2203.1 | 6156.6* | 2287.9# |
| **Miscellaneous** | | | | | | | | | | | | |
| Alb | 1420999.2 | 1613066.5 | 1253648.3# | 970715.9 | 1306413.8 | 871189.9 | 1423566.8 | 1005559.0* | 832522.3* | 1511958.1 | 1002958.3* | 805777.0*# |
| Hgf | 52.0 | 151.4* | 175.8* | 105.3 | 147.4 | 195.5 | 103.6 | 201.7* | 240.7* | 191.8 | 380.8* | 372.5* |

**Index of Genes**

List of genes in Supplemental Table 2

**Abca1**: ATP binding cassette subfamily A member 1, **Abcg5**: ATP-binding cassette sub-family G member 5, **Abcg8**: ATP-binding cassette sub-family G member 8, **Adam17**: A disintegrin and metalloprotease 17 (also known as Tace), **Adgre1**: adhesion G protein-coupled receptor E1 (also known as F4/80), **Aif1**: allograft inflammatory factor 1 (also known as Iba1), **Alb**: Albumin, **Casp6**: Caspase-6, **Cd68**: Cluster of Differentiation 68, **Col1a1**: Collagen, type 1 alpha 1, **Col1a2**: Collagen, type 1 alpha 2, **Col3a1**: Collagen, type 3 alpha 1, **Col5a2**: Collagen, type 5 alpha 2, **Csf1**: Colony stimulating factor 1, **Cyp8b1**: Cytochrome P450, family 8, subfamily B, polypeptide 1, **Fabp1**: Fatty Acid-Binding Protein 1, **Hgf**: Hepatocyte growth factor, **Hmgcr:** 3-hydroxy-3-methyl-glutaryl-coenzyme A reductase, **Hmgcs1**: 3-hydroxy-3-methylglutaryl-Coenzyme A synthase 1, **Insig1**: Insulin induced gene 1, **Itgam**: Integrin alpha M, **Ldlr**: Low-density lipoprotein receptor, **Lgals3**: Galectin-3, **Lpin1**: Lipin-1, **Mmp12**: Matrix metalloproteinase-12, **Pcsk9**: Proprotein convertase subtilisin/kexin type 9, **Plin1**: Perilipin-1, **Plin2**: Perilipin-2, **Plin3**: Perilipin-3, **Plin4**: Perilipin-4, **Plin5**: Perilipin-5, **Saa3**: Serum amyloid A3, **Scap**: Sterol regulatory element-binding protein cleavage-activating protein, **Soat1**: Sterol O-acyltransferase, **Sqle**: Squalene monooxygenase, **Trem2**: Triggering receptor expressed on myeloid cells 2, **Tyrobp**: TYRO protein tyrosine kinase-binding protein.

**Supplemental table 3.** Comparison of C57BL/6J mice injected with AAV-sTREM2 versus AAV-control and then assigned to HFHC diet for 15 and 90 days for physical attributes, blood chemistry, and histology.

| | 15 Days | | 90 days | |
|---|---|---|---|---|
| | **AAV-control** n=6 | **AAV-sTREM2** n=6 | **AAV-control** n=10 | **AAV-sTREM2** n=10 |
| Body weight, *g*, mean (SD) | 37.9 (1.8) | 40.3 (3.1) | 45.4 (4.1) | 45.8 (5.3) |
| Liver Weight, *g*, mean (SD) | 1.7 (0.2) | 1.9 (0.3) | 3.5 (1.0) | 3.7 (1.1) |
| Liver/Body weight ratio, *%*, mean (SD) | 4.5 (0.3) | 4.8 (0.5) | 7.7 (1.7) | 8.0 (1.7) |
| Food Intake, *g/mouse/day*, mean (SD) | 2.7 (0.2) | 2.9 (0.2) | 3.2 (0.3) | 3.2 (0.2) |
| Calories Intake, *Kcal/ mouse/day*, mean (SD) | 11.2 (0.8) | 12.0 (0.83) | 13.3 (1.2) | 13.3 (0.8) |
| ALT, *U/L*, mean (SD) | 37.8 (7.6) | 58.2 (24.8) | 193.2 (72.2) | 317.6 (219.3) |
| AST, *U/L*, mean (SD) | 49.5 (5.7) | 55.0 (8.9) | 134.6 (41.6) | 235.3 (177.9) |
| ALP, *U/L*, mean (SD) | 50.5 (4.5) | 59.0 (11.7) | 108.6 (23.4) | 134.4 (71.2) |
| Total Cholesterol, *mg/dL*, mean (SD) | 155.7 (14.1) | 167.8 (8.5) | 207.3 (51.4) | 222.4 (70.5) |
| HDL Cholesterol, *mg/dL*, mean (SD) | 96.8 (5.3) | 92.7 (6.8) | 104.0 (23.6) | 109.0 (28.1) |
| LDL Cholesterol, *mg/dL*, mean (SD) | 9.8 (1.3) | 12.0* (0.6) | 15.6 (3.7) | 20.6 (7.8) |
| Glucose, *mg/dL*, mean (SD) | 326.0 (46.6) | 334.8 (54.2) | 327.7 (84.5) | 303.0 (96.4) |
| Triglycerides, *mg/dL*, mean (SD) | 60.0 (7.7) | 60.0 (10.5) | 58.8 (9.1) | 65.5 (13.7) |
| Steatosis grade, mean (SD) | 1.7 (0.6) | 2.3 (0.6) | 2.8 (0.5) | 3.0 (0.0) |
| Sirius red (Fibrosis) polarization at 200x magnification, *% area*, mean (SD) | 0.01 (0.01) | 0.01 (0.02) | 0.02 (0.01) | 0.09 (0.12) |
| Birefringence (crystals containing cholesterol), % area, mean (SD) | 0 | 0 | 0 | 0 |
| Trem2 staining at 200x magnification, *% area*, mean (SD) | 0.40 (0.28) | 26.85 (14.29)** | 0.15 (0.14) | 3.17 (3.28)** |
| Cd68 staining at 200x magnification, *% area*, mean (SD) | 1.41 (0.44) | 1.30 (0.36) | 1.50 (0.73) | 1.38 (0.72) |

*Indicates that the comparison of AAV-Trem2 vs AAV-Control was statistically significant (*p<0.05; **p<0.01)

**Supplemental figure 1.** The structure of the AAV-control and AAV-sTREM2 vectors. An AAV-DJ/8 encoding for the extra-cellular shedded portion (soluble) of Trem2 under the control of the liver-specific human thyroxine binding globulin promoter (AAV-Trem2) was generated. The same AAV-DJ/8 without the sTrem2 sequence (AAV-control) was used as a control. The vectors were generated by the CRISPR, Vector and Transgenic Mouse Core at the University of Washington's South Lake Union Campus.

Empty TBG-AAV8

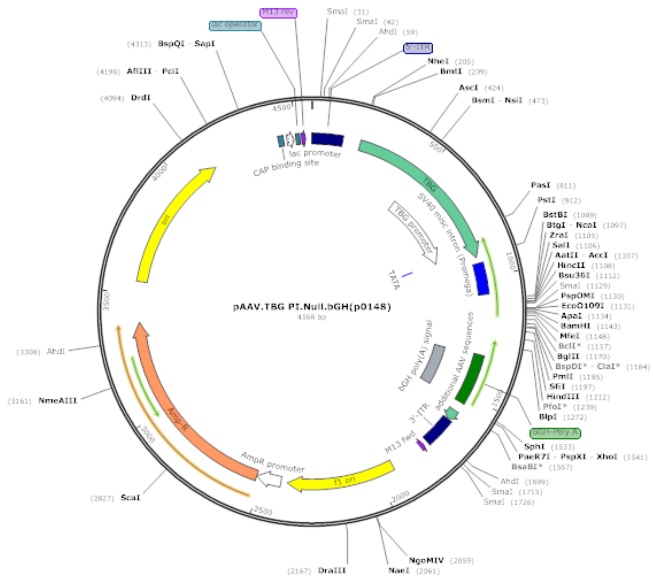

TBG-AAV8 with sTrem2 sequence

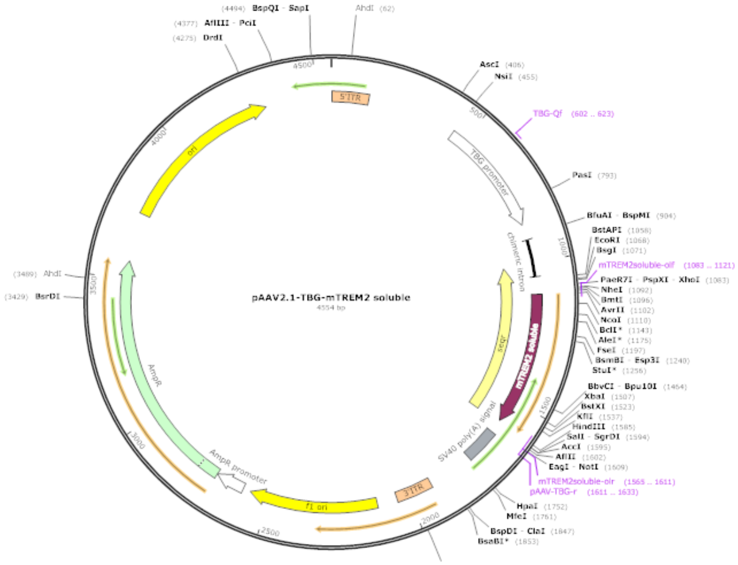

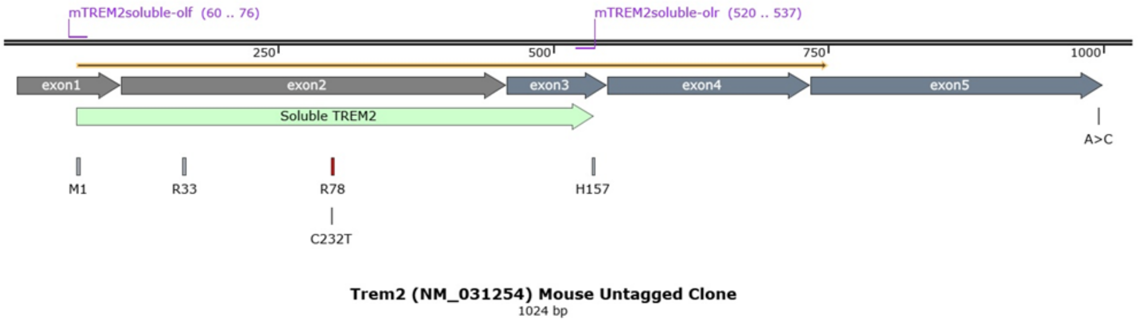

**Supplemental figure 2.** Comparison of C57BL/6J mice assigned to normal diet, HF diet and HFHC diet for 90 days. Livers from mice fed a normal, HF, or HFHC diet for 90 days were stained with H&E and Sirius Red. Samples were imaged at 200x magnification under brightfield and polarized light. Although there was substantial macrovesicular steatosis in the HF and HFHC-fed mice after 90 days on the assigned diets, there was no evidence of fibrosis by Sirius red staining and no evidence of birefringence under polarized light (that would be indicative of cholesterol crystallization).

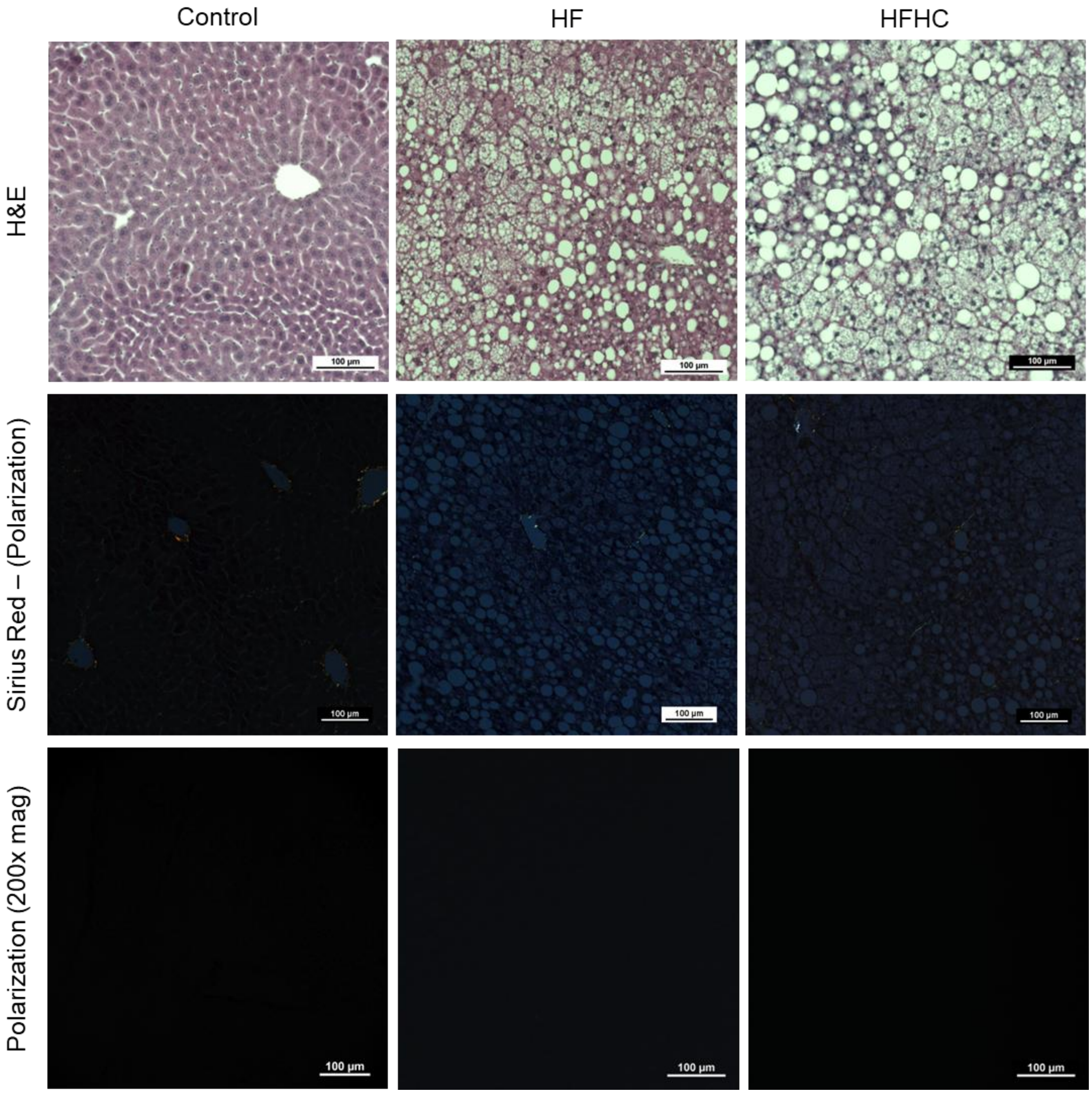
